# Neuronal quantification in the primary motor cortex of mouse brains fixed with solutions from human gross anatomy laboratories

**DOI:** 10.64898/2026.08.24.744656

**Authors:** Amy Gérin-Lajoie, Eve-Marie Frigon, Walter Adame-Gonzalez, Mahsa Dadar, Denis Boire, Josefina Maranzano

**Affiliations:** Department of Anatomy, University of Québec in Trois-Rivières, Trois-Rivières, Quebec, Canada; Department of Psychiatry, McGill University, Montreal, Quebec, Canada; Douglas Mental Health University Institute, Montreal, Quebec, Canada; Department of Neurology and Neurosurgery, Montreal Neurological Institute, McGill University, Montreal, Quebec, Canada

**Author notes:** Corresponding author: Amy Gérin-Lajoie, Université du Québec à Trois-Rivières, 3351 Bd des Forges, Trois-Rivières, Québec, Canada, G8Z 4M3.

**Keywords:** neurons, primary motor cortex, segmentation, quantification, immunohistochemistry

## Abstract

**Background:** Brain banks usually provide small tissue blocks fixed by immersion in neutral-buffered formalin (NBF). While still underexploited for research, gross anatomy laboratories could provide full brains fixed by perfusion with solutions better suited for gross anatomy dissection. However, the chemicals in these solutions might have a different impact on histology protocols for cell quantification than in NBF-fixed brains. The main goal of this study is to compare the effects on the number and size of labeled neurons of the primary motor cortex (PMC) of mouse brains fixed with three different solutions: (1) NBF, typical of brain banks, (2) a saturated salt solution (SSS), and (3) an alcohol-formaldehyde solution (AFS), both used in human anatomy laboratories.

**Methods:** 27 C57BL/6J mouse brains were perfused with the NBF (N=9), SSS (N=9) or AFS (N=9), then cut in 40-μm slices and processed with immunohistochemistry to target neurons. Various quantitative variables were assessed manually and automatically on photomicrographs of 3 regions of interest (ROIs) of the PMC per specimen, namely the total and individual neuronal profile areas, number and diameters. The effects of the three fixatives on these variables were compared using ANOVA or Kruskal-Wallis, depending on the distribution. For measures on individual cells, a generalized linear mixed model was applied. Dice coefficients and correlations were applied to evaluate the agreement of the manual and automatic methods.

**Results:** There was no significant difference between the brains fixed by the three fixatives for the total and individual cell areas, the total cell count and the cell diameters. The values obtained from manual and automatic measures had an overall good agreement (Dice coefficients > 0.79).

**Conclusion:** It was found that the SSS and AFS had similar impacts on the quantitative variables in the tissue as the NBF. These results are promising for neuroscientists interested in using brains from anatomy laboratories for quantitative research on neurons from the PMC.

## 1. Introduction

In neuroscientific research, histology remains the gold standard to assess the cellular and molecular changes caused by normal and pathological aging of the brain at the microstructural level (den Bakker, 2017; Frigon et al., 2022b; Frigon et al., 2024; Frigon et al., 2026; Hoffmann et al., 2011; Ma et al., 2024). Since biopsies are an invasive and rarely performed technique to acquire brain tissue for histology processing, researchers tend to turn to *ex vivo* approaches, which allow them to gather information concerning the cellular changes. *Ex vivo* specimens need to be fixed to prevent tissue decomposition and to preserve the cellular architecture (Arnold et al., 1996; Brenner, 2014; Musiał et al., 2016; Thavarajah et al., 2012). Fixation involves the use of chemical solutions that could potentially modify the tissue’s physicochemical characteristics (Holmes et al., 2017). The solutions might in turn alter the distribution of fluids and the molecular conformation of proteins, which translates into deformations and changes in cellular morphology (Alkemade et al., 2023; Haga et al., 2019; Kotrotsou et al., 2014; Vasung et al., 2019).

Neutral-buffered formalin solution (NBF) remains the most widely used chemical fixative in neuroscientific and histopathological studies. It prevents tissue decay through protein cross-linking (Blum, 1896; de Paula et al., 2018; Grigorev & Korzhevskii, 2018; Helander, 1994; Hobro & Smith, 2017; Hopwood, 1985; Ma et al., 2024; O’Rourke & Padula, 2016; Puchtler & Meloan, 1985; Wu et al., 2022). Although formaldehyde has excellent tissue preservation properties that prevent decomposition very effectively (Fox & Benton, 1987; O’Rourke & Padula, 2016), it also causes tissue shrinkage and hardening (Balta et al., 2015; Barton et al., 2009; Birkl et al., 2016; Brenner, 2014; Eisma et al., 2013; Eisma et al., 2011; Frigon et al., 2022b; Hayashi et al., 2014; Hayashi et al., 2016; Jaung et al., 2011; Mayer, 2006; Regelsberger et al., 2015; Richins et al., 1963; Tomalty et al., 2019; Weisbecker, 2012). Furthermore, it is a toxic compound with carcinogenic potential in high concentrations and with long-term exposure (Benet et al., 2014; Brenner, 2014; Coleman & Kogan, 1998; de Paula et al., 2018; Fischer, 1905; Hayashi et al., 2016; Lombardero et al., 2017; Moelans et al., 2011; Musiał et al., 2016; Otsuka et al., 2022). Consequently, specimens fixed exclusively with NBF are not appropriate for gross anatomy laboratories. Alternative solutions have been developed and optimized to better mitigate these challenges for the dissection and teaching of gross anatomy. These solutions include a saturated salt solution (SSS) (Coleman & Kogan, 1998; Hayashi et al., 2014; Hayashi et al., 2016) and an alcohol-formaldehyde solution (AFS) (Benet et al., 2014). Both solutions contain lower formaldehyde concentrations, combined with other chemicals (i.e., sodium chloride and various types of alcohols) to allow for lengthy dissections and to decrease the health hazards while maintaining an excellent tissue preservation.

Although the SSS and AFS are useful solutions for gross anatomy laboratories, their compatibility with (immuno-) histochemical procedures have not been extensively tested. We have previously shown that they appropriately preserved antigenicity of the four main brain cell populations (i.e., neurons, astrocytes, microglia, and myelin of oligodendrocytes) in mice (Frigon et al., 2022b) and humans (Frigon et al., 2024). These studies revealed satisfactory tissue quality of the AFS-fixed samples, but lower tissue quality and ease of manipulation of the sections of tissue fixed with the SSS. The results also highlighted that SSS-fixed cells always had irregular, shriveled cell contours. However, since both studies focused on qualitative assessments, whether these solutions impact quantitative studies of brain tissue has yet to be determined.

Indeed, it is widely known that neuronal loss is a hallmark of many neurodegenerative diseases and other neurological pathologies (Agnello & Ciaccio, 2022; Vogels et al., 1990). Morphological alterations are also common. For instance, neuronal shrinkage within the nucleus basalis of Meynert has been detected in cases of Alzheimer’s disease (Rinne et al., 1987; Vogels et al., 1990). It has also been observed in the hippocampus, substantia nigra and locus coeruleus of brains of patients with schizophrenia (Rajkowska et al., 1998). The number and size of pyramidal cells within the primary motor cortex (PMC) of people with amyotrophic lateral sclerosis are also known to be impaired (Hammer et al., 1979; Mezzapesa et al., 2013). Consequently, it is important to explore the neuronal count, size and area in brains fixed with each fixative to elucidate whether solutions used in anatomy laboratories affect the cell populations in the same way as the NBF used by brain banks. In doing so, we aim to support the use of an alternative source of brain samples for research purposes and neuropathological studies, since brain banks generally only provide small blocks of NBF-fixed tissue while gross anatomy laboratories could offer full brains.

Therefore, the present study aims to describe and compare the effects of two solutions used in gross anatomy laboratories (i.e., the SSS and AFS) to those of the classic fixative used by brain banks (i.e., the NBF) on the number and size of neurons of the PMC, using manual and automatic techniques. The study was first realized on mice to avoid the confounding variables associated with human tissue. We hypothesized that the cell number, profile area and diameters may be different in mouse brains fixed with the AFS and SSS compared to NBF-fixed brains. Since the formaldehyde concentration is higher in the NBF composition, we expected it to reduce the size and diameters of cells in comparison with the SSS and AFS, as formaldehyde is known to cause tissue shrinkage.

## 2. Materials and Methods

### 2.1 Population

We used a sample of 27 adult C57BL/6J mice (male to female ratio of 14 to 13) (Table 1) that were raised and kept in a controlled environment (45-60% humidity; 20-25°C temperature; 12-hour daylight schedule). We followed the guidelines as described by the Canadian Council on Animal Care. Our protocols were approved by the Ethics Committee of the University of Québec in Trois-Rivières.

**Table 1.**
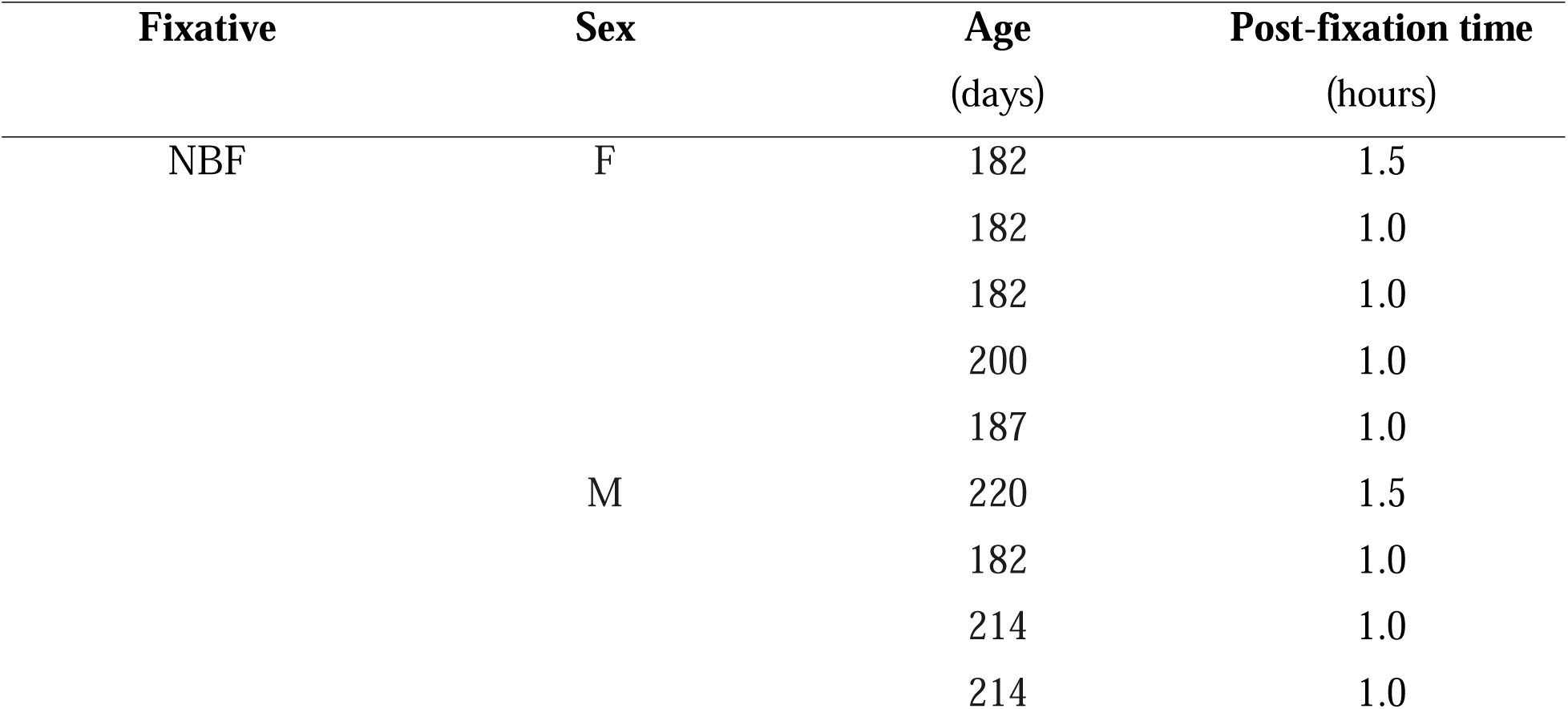
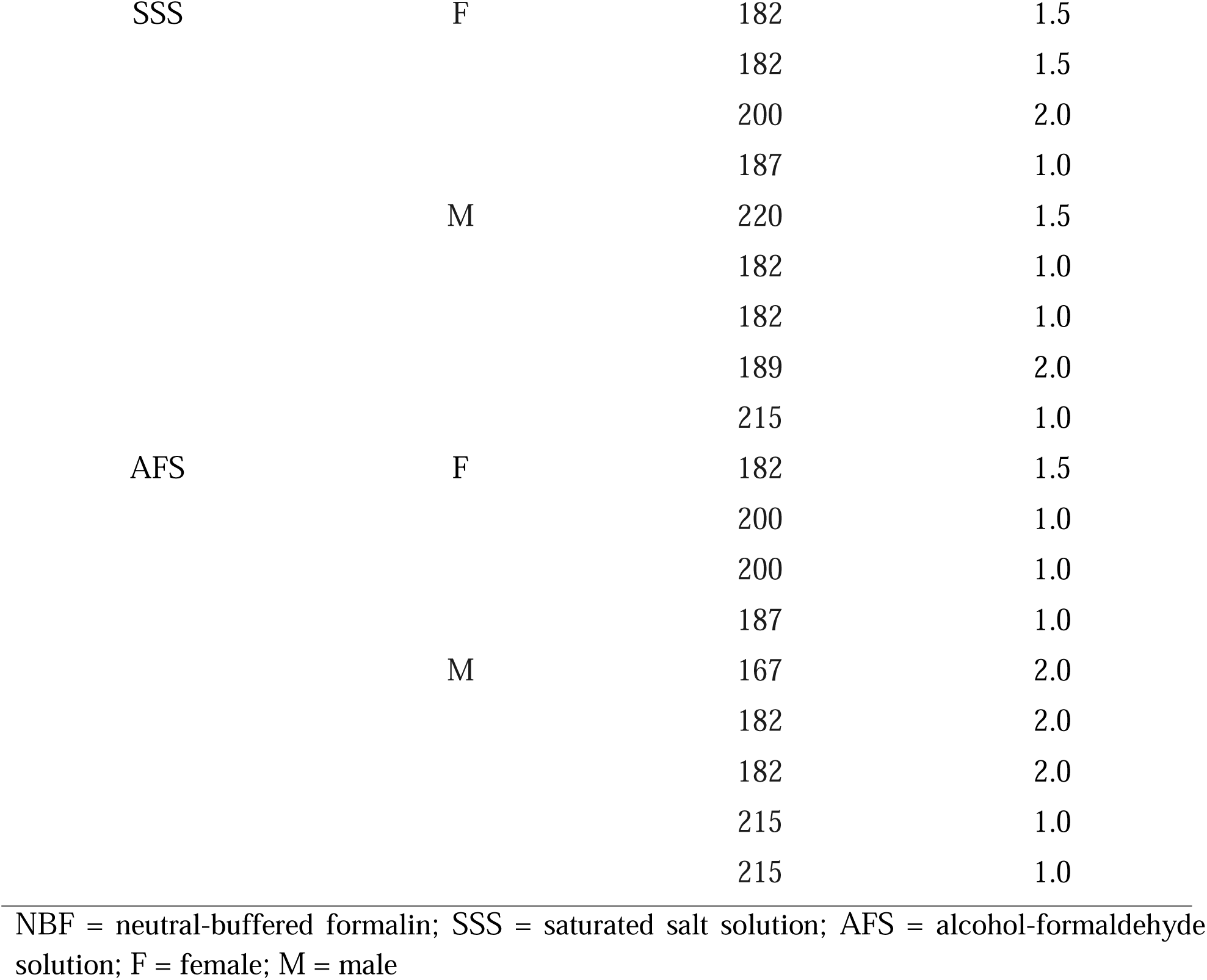
Experimental animals data.

By using mouse brains, we avoided the confounding variables associated with the use of human samples, such as variable post-mortem delay, fixation delay, comorbidities, biological heterogeneity of donors, etc.

### 2.2 Fixation procedure

Brains were fixed by transcardiac perfusion (Paul et al., 2008; Rana et al., 2022), using 0.1M phosphate-buffered saline (PBS) (0.9% NaCl). This was followed by the injection of one of the three fixative solutions tested in this study: 1) NBF, 2) SSS, and 3) AFS (see Table 2 for the chemical composition of each fixative) for 5 minutes to standardise fixative exposure. Mice were randomly assigned to fixative solution groups (NBF: N=9; SSS: N=9; AFS: N=9). Following the perfusion, the brains were harvested and immersed in their respective fixative solution for a post-fixation time before immunohistochemistry (IHC) processing. The length of the post-fixation was determined by assessing the color of the tissue, which indicates the amount of remaining blood (Frigon et al., 2022b): 2 hours for pink brains, 1 hour and a half for heterogeneous brains, and 1 hour for beige brains. The post-fixation times were not statistically different between the three groups (see Table 1).

**Table 2.** Chemical composition of the fixatives.

| **NBF** | **SSS** | **AFS**<br>(Benet et al., 2014) |
| --- | --- | --- |
| 4% formaldehyde | 0.8% formaldehyde | 2.3% formaldehyde |
| 0.1M PBS | 36% NaCl | 62.4% ethanol |
|  | 0.72% phenol | 10.2% phenol |
|  | 2% glycerol | 17% glycerol |
|  | 16% isopropyl alcohol |  |
|  | 2.5% Dettol |  |

NBF = neutral-buffered formalin; SSS = saturated salt solution; AFS = alcohol-formaldehyde solution; PBS = phosphate-buffered saline; NaCl = sodium chloride

### 2.3 Immunohistochemistry processing

After post-fixation, the brains were rinsed in 0.1 M PBS and immersed overnight in 30% sucrose for cryoprotection. Brains were then frozen in dry ice and stored at -80°C until processed. They were cut in 40-μm-thick coronal sections using a cryostat at -19°C (Leica CM1950). The sections were rinsed three times in five-minute intervals in 0.1M PBS before a 30-minute incubation in an aqueous methanol solution (20% methanol, 0.5% H_2_O_2_, and 0.3% Triton X-100 in 0.1M PBS) to quench endogenous peroxidase. The sections were rinsed three other times for five minutes in 0.1M PBS, followed by a two-hour incubation in a blocking solution comprising 3% normal donkey serum, 0.5% bovine serum albumin, and 0.3% Triton X-100 in 0.1M PBS. The sections were then transferred directly into the same blocking solution with the added primary antibodies (1:3000, rabbit anti-neuronal nuclei (NeuN), Abcam, Ab177487) for an overnight incubation at 4°C on a shaker. The next day, the slices were rinsed three times at five-minute intervals in 0.1M PBS before an incubation in the blocking solution with secondary antibodies (1:500, donkey anti-rabbit biotinylated, NovusBio, NBP1-75274) for two hours at room temperature. The sections were then rinsed again in 0.1M PBS in three five-minute baths before a 30-minute incubation in the dark in an avidin-biotin complex kit (Vector Laboratories, Newar, CA USA, catalog #VECTPK6100). Following three five-minute rinses in 0.05M TRIS-buffered saline (TBS) (0.9% NaCl), sections were then incubated for ten minutes in 0.07% diaminobenzidine (MilliporeSigma, Burlington, MA, USA catalog #D5905-50TAB) with 0.024% H_2_O_2_ in 0.05M TBS. The sections were rinsed three times for five minutes in 0.1M phosphate buffer and mounted on 2% gelatin-subbed slides. Finally, the slides were dehydrated in graded alcohols (70% ethanol, 95% ethanol, 2 x 100% ethanol, and 2 x xylene) and coverslipped with Eukitt quick-hardening mounting medium.

### 2.4 Microscopy

Photomicrographs were captured through a brightfield microscope (Olympus, Tokyo, Japan, BX51W1) controlled by Neurolucida (MBF Bioscience, Williston, VT, USA) with a 100X objective (100X, UPlanSApo 100x/1.40 Oil∞/0.17/FN26.5) and a high resolution DV-HR-CLR Lumina camera with a 1”CCS sensor. The boundaries of the PMC were traced on the Neurolucida software from figure 33 (0.22 mm posterior to Bregma) of the atlas of Paxinos and Franklin (2001). Using cytoarchitectural landmarks, three regions of interest (ROIs) (125 x 100 microns) were selected in layer 3 of the PMC. Three images were captured for each selected ROI across the thickness of the tissue section, including one at the top, one in the middle and one at the bottom of the slice (hereby referred to as top, middle, and bottom, respectively). An example of the delimitations of the histological sections is shown in Figure 1.

**Figure 1.**
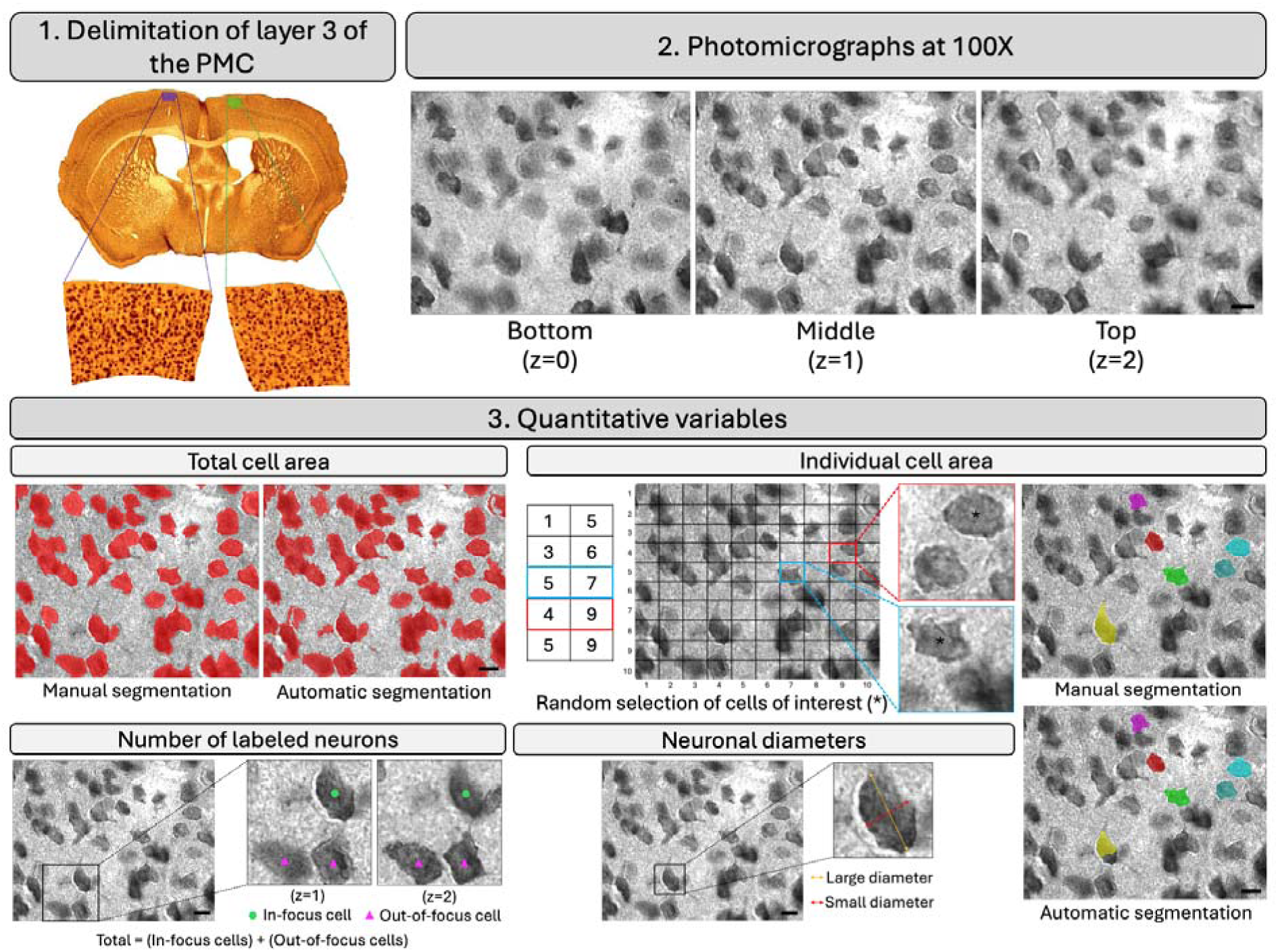
Summary of the image analyses. PMC = primary motor cortex; z = level of thickness of the histological section. Scalebar = 10 μm.

### 2.5 Image analyses

Quantification of the total neuronal profile area, the area of 10 individual neurons, the total neuron count, and their diameters were considered for subsequent statistical analyses. All variables are illustrated in Figure 1.

#### 2.5.1 Total cell area

The total neuronal area was defined as the sum of all the neuronal profile areas within each photomicrograph. This variable is therefore a composite of the number of cells within a ROI and their individual sizes.

##### 2.5.1.1 Manual segmentation

The total cell area was segmented manually by the same rater (A.G.L.) in the middle image of each ROI using the Display software (MINC Tool Kit, McConnell BIC). An intensity threshold function allowed us to select the range of gray intensity of pixels that best revealed neuronal area on each image. All the pixels within the chosen range of intensities were selected by the software and then corrected manually to add false negative pixels and remove false positive pixels. These segmentations were used as the gold standard for the remainder of the project.

##### 2.5.1.2 Automatic segmentation

Using the same images, we also tested an automatic detection method that generated labels of the total cell area for every specimen. The method itself was based on an adapted in-house deep learning network for medical image segmentation using a UNet architecture (Ronnerberger et al., 2015). Raw MINC volumes were paired with their painted cell masks, and the masks were binarized with a > 0.5 threshold. Out of the 79 available samples (pixel dimensions of 3 x 2 192 x 2 752), 67 were used for training the network using five-fold cross-validation, leaving the remaining twelve reserved for the test subset. The images were fed into the network after linear min-max normalization to [-1, 1]. Due to the relatively small number of training images, aggressive augmentation techniques were applied to the input images during training: horizontal and vertical flips, random 90-degree rotations, additive white Gaussian noise (σ = 0.01), random gamma [0.8, 1.2], and CutMix (p = 0.5, min_frac_ = 0.05, max_frac_ = 0.2). For the validation and test subsets, no augmentations were used but they were processed with the same normalization to [-1, 1]. The network was trained using a standard implementation of binary cross-entropy (*BCELoss*) for 200 epochs per fold using a batch size of 1, and the checkpoint with the best validation loss per fold was used for downstream inference.

#### 2.5.2 Individual cell area

This variable allows for the evaluation of the effect of the fixative solution on cell areas, without the influence of cellular overlap or the confounding effects of other variables. In combination with the total cell area and the number of neurons, this metric allows to test whether the fixation has an influence on the neuropil or the cells, or on both.

##### 2.5.2.1 Manual segmentation

Ten individual cells were randomly selected per ROI with a 10 x 10 grid and a two-column random number generator (RANDOM.ORG, 2026). To be considered for inclusion in the sampling, cell profiles were required to be either isolated from other cells, or their borders needed to be very distinct from neighboring cells. This ensured that it could be segmented without inadvertently including portions of the neighboring cells. Then, every randomly selected cell was segmented manually (A.G.L.) using the same manual method used for the total cell area quantification. The rater measured ten cell areas per ROI and three ROIs per specimen, producing individual areas for 30 cells per specimen.

##### 2.5.2.2 Automatic segmentation

The automatic total cell areas were used as a basis to adapt the automatic segmentation for individual cell areas. The cell clusters were isolated using a watershed algorithm using the *scipy.ndimage.watershed_ift* Python package (distance_min_ = 90 px, distance_sigma_ = 0 px, area_min_ = 15 000 px, connectivity = 8). This permits the separation of overlapping objects that have previously been segmented by another method (Kornilov et al., 2022). The automatic area measures of the same cells of interest were then obtained as they were with the manual method. Any measure over twice the standard deviation was deemed an outlier and visually reassessed. The cells that were incorrectly or excessively segmented by false identification of surrounding neuropil and adjacent cells were removed before the statistical analysis (see Supplementary Figure 1 and Supplementary Table 1).

#### 2.5.3 Total cell count

This variable allows us to assess whether the fixative solution impacts the number of labeled neurons in a specific ROI. Either this variable can be affected by the changes in volume of the tissue, or by the effect of the chemicals present in the fixative solutions on antigenicity preservation.

The total number of labeled neurons per ROI was quantified by manually adding markers on each visible neuron using the Neurolucida software. Two different markers were used: one for in-focus cells and one for out-of-focus cells. We defined in-focus cells as cells with clear outlines in the middle layer (z = 1) and out-of-focus cells as cells with blurry outlines in the middle layer (z = 1) and clear outlines either in the bottom or top layer (z = 0 and z = 2). The purpose of the in-focus and out-of-focus cells is to consider all cells present in the middle layer while still differentiating their spatial characteristics, since they could be related to the neuronal size. Larger neurons would more often appear in focus because they have a higher probability of being included in the middle layer. Conversely, smaller neurons would be less likely to appear as often in the middle layer, as they would be more dispersed in the thickness of the slice. Finally, we also added up the number of in-focus and out-of-focus cells to obtain a total cell count.

#### 2.5.4 Cell diameters

The large and small diameters are informative of the size of individual cells. It is also the standard way in which biology literature has traditionally reported cell size (Paxinos & Mai, 2004). Furthermore, the diameters enable the assessment of cell morphology by calculating a circularity ratio.

The large and small diameters of the same 10 cells per ROI were manually measured using the Neurolucida software. The large diameter was defined as the length between the two furthest opposing points of the cell and the small diameter as the length between the two furthest opposing points perpendicular to the large diameter. The diameters were also measured automatically based on the previously generated automatic cell areas.

Using these cell diameters, a circularity ratio was measured by dividing the small diameters by the large diameters:

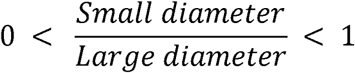

A value closer to one would mean the cell has a circular morphology, wherein the large and small diameters are similar in length. On the contrary, a value further from 1 would mean the cell had an elongated morphology, wherein the large diameters had a distinctly longer length than the small diameter. These ratios were measured for every cell and compared between solutions. They were generated for both the manually and automatically measured cell diameters.

### 2.6 Statistical analyses

One-way ANOVAs were performed when the distribution of values met the normality requirement according to the Kolmogorov-Smirnov test. In such scenarios, the mean and standard deviation were reported. Since the comparisons were made between three groups of equal size, Fisher’s Least Significant Difference (LSD) was applied as a post hoc test. Kruskal-Wallis tests were performed when the distribution of values did not respect the normality curve. In these cases, the median and range were instead reported.

For variables with multiple measures per specimen (e.g., individual cell area, cell diameters, circularity ratios), a generalized linear mixed model (GLMM) was performed to account for the grouped values. The formula was the following:

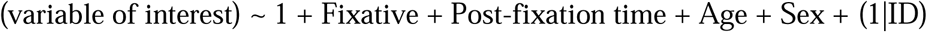

The validity and reproducibility of the individual cell manual segmentation was assessed by evaluating the intraclass correlation coefficients (ICCs) for intra-rater (A.G.L.-1 vs A.G.L.-2) and inter-rater (A.G.L. vs J.M.) variabilities. A two-way mixed ICC model for absolute agreement was selected to evaluate the agreement across raters.

The comparisons between manual and automatic segmentation methods were done with the Dice coefficient. All comparisons between manual and automatic measures were also evaluated with correlations. Both the Pearson and Spearman correlations were used, depending on the data normality.

All statistical tests were performed using SPSS (29.0.1.1), except for the GLMMs that were performed with MATLAB (R2025b). Violin plots were generated with GraphPad Prism (10.4.1).

## 3. Results

To facilitate the comparison of results between the fixatives, all descriptive statistics, Dice coefficients and correlations are summarized in Table 3. The results of ANOVAs or Kruskal-Wallis tests as well as GLMMs are detailed in their respective sections for each variable.

**Table 3.** Summary of the descriptive statistics for each fixative.

|  |  | NBF |  | SSS |  | AFS |  |
| --- | --- | --- | --- | --- | --- | --- | --- |
|  |  | Manual | Automatic | Manual | Automatic | Manual | Automatic |
| Total cell area | Mean | 4 836 | 5 240 | 5 268 | 5 506 | 5 032 | 5 216 |
|  | SD | 1 164 | 607 | 910 | 658 | 706 | 574 |
|  | Median | 4 991 | 5 200 | 5 263 | 5 651 | 5 093 | 5 358 |
|  | Range | 2 505 - 6 526 | 4 148 - 5 945 | 3 928 - 6 432 | 4 442 - 6 252 | 3 881 - 5 979 | 4 176 - 6 014 |
|  | Dice coefficient | 0.89 |  | 0.93 |  | 0.95 |  |
|  |  | 0.92 |  |  |  |  |  |
|  | Pearson correlation | r = 0.849; p = 0.004 |  | r = 0.904; p < 0.001 |  | r = 0.911; p < 0.001 |  |
|  |  | r = 0.867; p < 0.001 |  |  |  |  |  |
| Individual cell area | Mean | 77.2 | 101.7 | 81.0 | 109.1 | 79.7 | 99.8 |
|  | SD | 23.0 | 47.6 | 24.0 | 50.5 | 24.7 | 44.8 |
|  | Median | 73.3 | 87.6 | 78.2 | 95.2 | 74.8 | 86.1 |
|  | Range | 35.0 - 158.0 | 36.0 - 255.0 | 27.0 - 189.0 | 34.0 - 259.0 | 26.0 - 164.0 | 32.9 - 255.0 |
|  | Dice coefficient | 0.79 |  | 0.81 |  | 0.83 |  |
|  |  | 0.81 |  |  |  |  |  |
|  | Spearman correlation | r<sub>s</sub> = 0.392; p < 0.001 |  | r<sub>s</sub> = 0.507; p < 0.001 |  | r<sub>s</sub> = 0.599; p < 0.001 |  |
|  |  | r<sub>s</sub> = 0.502; p < 0.001 |  |  |  |  |  |
| Total cell count | Mean | 88 | NA | 80 | NA | 83 | NA |
|  | SD | 19 |  | 13 |  | 12 |  |
|  | Median | 84 |  | 78 |  | 82 |  |
|  | Range | 58 - 128 |  | 57 - 102 |  | 60 - 101 |  |
| In-focus cell count | Mean | 24 | NA | 30 | NA | 33 | NA |
|  | SD | 7 |  | 5 |  | 6 |  |
|  | Median | 25 |  | 30 |  | 35 |  |
|  | Range | 12 - 35 |  | 24 - 37 |  | 21 - 40 |  |
| Out-of-focus cell count | Mean | 64 | NA | 50 | NA | 50 | NA |
|  | SD | 17 |  | 9 |  | 9 |  |
|  | Median | 65 |  | 50 |  | 51 |  |
|  | Range | 29 - 93 |  | 33 - 65 |  | 39 - 62 |  |
| Large diameters | Mean | 11.86 | 14.20 | 12.30 | 14.53 | 12.35 | 14.34 |
|  | SD | 2.20 | 3.85 | 2.21 | 4.10 | 2.62 | 3.98 |
|  | Median | 11.56 | 13.27 | 11.97 | 13.49 | 11.96 | 13.23 |
|  | Range | 7.73 - 22.49 | 7.71 - 24.23 | 6.83 - 22.38 | 7.24 - 24.28 | 5.99 - 24.75 | 7.81 - 23.75 |
|  | Spearman correlation | r<sub>s</sub> = 0.313; p < 0.001 |  | r<sub>s</sub> = 0.453; p < 0.001 |  | r<sub>s</sub> = 0.503; p < 0.001 |  |
|  |  | r<sub>s</sub> = 0.423; p < 0.001 |  |  |  |  |  |
| Small diameters | Mean | 9.02 | 9.75 | 9.18 | 10.15 | 9.01 | 9.49 |
|  | SD | 1.59 | 2.20 | 1.80 | 2.17 | 1.68 | 2.07 |
|  | Median | 9.06 | 9.41 | 9.02 | 9.86 | 9.06 | 9.15 |
|  | Range | 4.74 - 14.58 | 5.36 - 15.91 | 4.54 - 15.96 | 5.14 - 16.10 | 5.05 - 14.95 | 4.93 - 15.54 |
|  | Spearman correlation | r<sub>s</sub> = 0.373; p < 0.001 |  | r<sub>s</sub> = 0.473; p < 0.001 |  | r<sub>s</sub> = 0.565; p < 0.001 |  |
|  |  | r<sub>s</sub> = 0.469; p < 0.001 |  |  |  |  |  |
| Circularity ratios | Mean | 0.77 | 0.71 | 0.76 | 0.73 | 0.75 | 0.69 |
|  | SD | 0.13 | 0.12 | 0.14 | 0.13 | 0.14 | 0.14 |
|  | Median | 0.77 | 0.72 | 0.75 | 0.74 | 0.77 | 0.69 |
|  | Range | 0.44 - 1.00 | 0.38 - 0.99 | 0.39 - 0.99 | 0.35 - 1.00 | 0.38 - 0.99 | 0.36 - 0.99 |
|  | Spearman correlation | r<sub>s</sub> = 0.326; p < 0.001 |  | r<sub>s</sub> = 0.452; p < 0.001 |  | r<sub>s</sub> = 0.547; p < 0.001 |  |
|  |  | r<sub>s</sub> = 0.444; p < 0.001 |  |  |  |  |  |

Areas are reported in square microns. Diameters are reported in microns. NBF = neutral-buffered formalin; SSS = saturated salt solution; AFS = alcohol-formaldehyde solution; SD = standard deviation; $r$ = Pearson correlation coefficient; $r_s$ = Spearman correlation coefficient

### 3.1 Total cell area

Using the manual segmentation method, the mean total cell area was 4 836 ± 1 164 μm^2^ in NBF-fixed specimens, 5 268 ± 910 μm^2^ in SSS-fixed specimens, and 5 032 ± 706 μm^2^ in AFS-fixed specimens. The manually measured total cell area was not significantly different between the three fixative groups (ANOVA: F_2;24_ = 0.471; p = 0.630; η^2^_p_ = 0.038) (Figure 2A).

**Figure 2.**
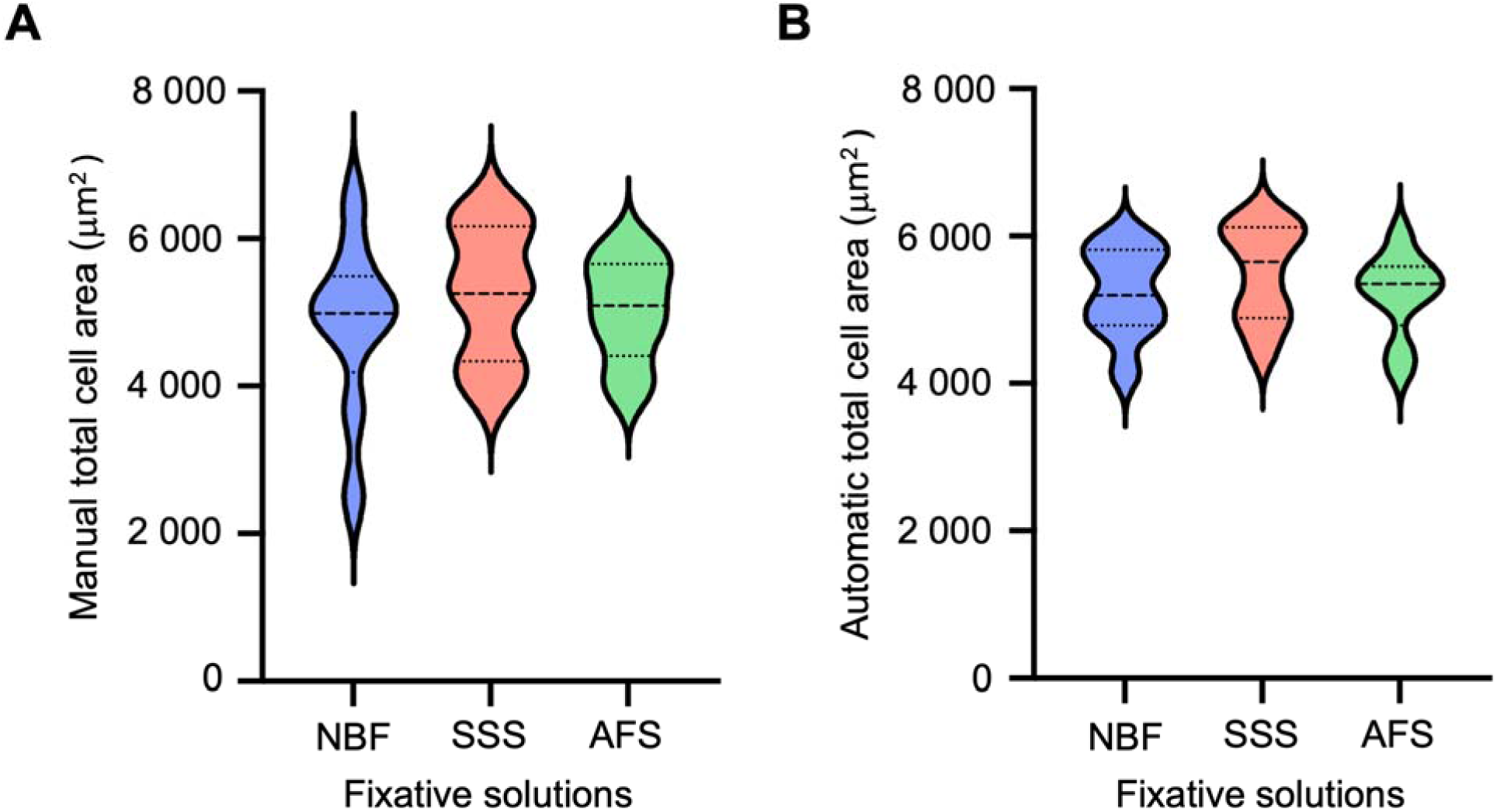
Violin plots of the total cell areas in all three fixative solutions. A) Manually segmented total cell areas. B) Automatically segmented total cell areas. μm = micrometers; NBF = neutral-buffered formalin; SSS = saturated salt solution; AFS = alcohol-formaldehyde solution

Using the automatic segmentation method, the mean total cell area was 5 240 ± 607 μm^2^ in NBF-fixed specimens, 5 506 ± 658 μm^2^ in SSS-fixed specimens, and 5 216 ± 574 μm^2^ in AFS-fixed specimens (Figure 2B).

Across all three fixatives, the Dice coefficient between the manual and automatic total cell areas showed a pixel alignment of 0.92. The Dice coefficient was 0.89 for NBF-fixed specimens, 0.93 for SSS-fixed specimens, and 0.95 for AFS-fixed specimens. There was a significant Pearson correlation between the manual and automatic total cell areas (r = 0.867; p < 0.001). Within each fixative group, these correlations were significant for NBF-fixed specimens (r = 0.849; p = 0.004), SSS-fixed specimens (r = 0.904; p < 0.001), and AFS-fixed specimens (r = 0.911; p < 0.001).

### 3.2 Individual cell area

Using the manual segmentation, the median individual cell area was 73.3 μm^2^ (range: 35.0-158.0 μm^2^) in NBF-fixed specimens, 78.2 μm^2^ (range: 27.0-189.0 μm^2^) in SSS-fixed specimens, and 74.8 μm^2^ (range: 26.0-164.0 μm^2^) in AFS-fixed specimens. The manually segmented individual cell area was not significantly different between NBF- and SSS-fixed specimens (GLMM: *t*(804) = 0.576; p = 0.565) or between NBF- and AFS-fixed specimens (GLMM: *t*(804) = -0.218; p = 0.827) (Figure 3A).

**Figure 3.**
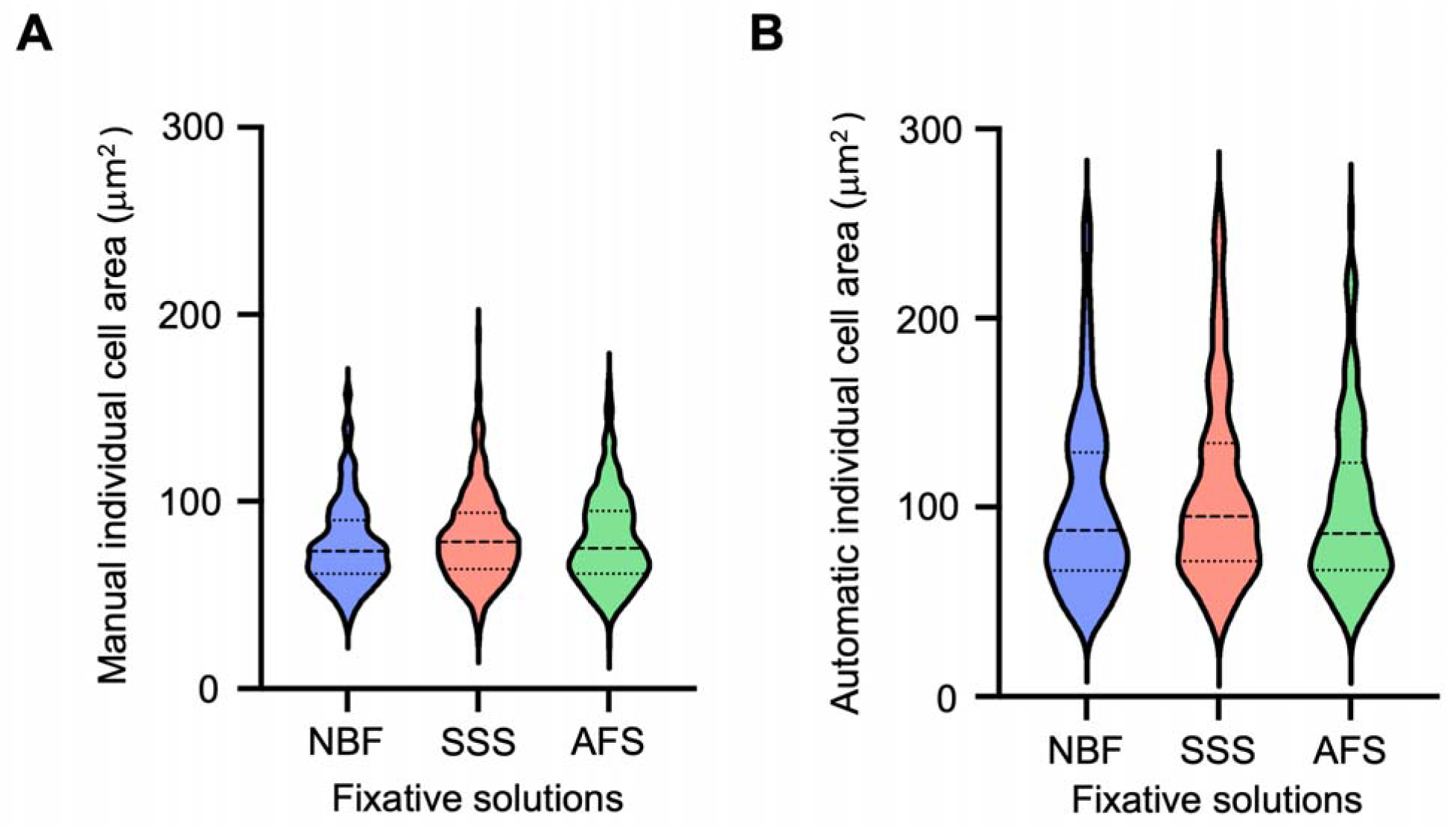
Violin plots of the individual cell areas in all three fixative solutions. A) Manually segmented individual cell areas. B) Automatically segmented individual cell areas. μm = micrometers; NBF = neutral-buffered formalin; SSS = saturated salt solution; AFS = alcohol-formaldehyde solution

For the manual segmentation, the intra-rater variability (AGL-1 vs AGL-2; N=8) generated an ICC score of 0.889 [0.772-0.936], which shows a good to excellent level of reliability (Koo & Li, 2016). The inter-rater variability (mean of AGL-1 and AGL-2 vs JM; N=8) produced an ICC value of 0.686 [0.028-0.873], which shows an overall moderate reliability (Koo & Li, 2016).

In contrast, using the automatic segmentation method, the median individual cell area was 87.6 μm^2^ (range: 36.0-255.0 μm^2^) in NBF-fixed specimens, 95.2 μm^2^ (range: 34.0-259.0 μm^2^) in SSS-fixed specimens, and 86.1 μm^2^ (range: 32.9-255.0 μm^2^) in AFS-fixed specimens (Figure 3B).

The mean Dice coefficient for all three groups showed a pixel-level alignment between the manual and automatic individual cells of 0.81. Per solution, the mean Dice coefficients were 0.79 for NBF-fixed specimens, 0.81 for SSS-fixed specimens, and 0.83 for AFS-fixed specimens. Moreover, there was a significant Spearman correlation between the manual and automatic individual cell areas (r_s_ = 0.502; p < 0.001). The correlations were also significant within each fixative group (r_s_ = 0.392; p < 0.001 for NBF-fixed specimens, r_s_ = 0.507; p < 0.001 for SSS-fixed specimens, and r_s_ = 0.599; p < 0.001 for AFS-fixed specimens).

### 3.3 Cell counts

#### 3.3.1 Total cell count

The mean total cell count per specimen was 87.56 ± 18.77 neurons in NBF-fixed specimens, 80.00 ± 13.39 neurons in SSS-fixed specimens, and 82.81 ± 12.30 neurons in AFS-fixed specimens. The total cell count was not significantly different between the three fixative groups (ANOVA: F_2;24_ = 131.269; p = 0.569; η^2^_p_ = 0.046) (Figure 4A).

**Figure 4.**
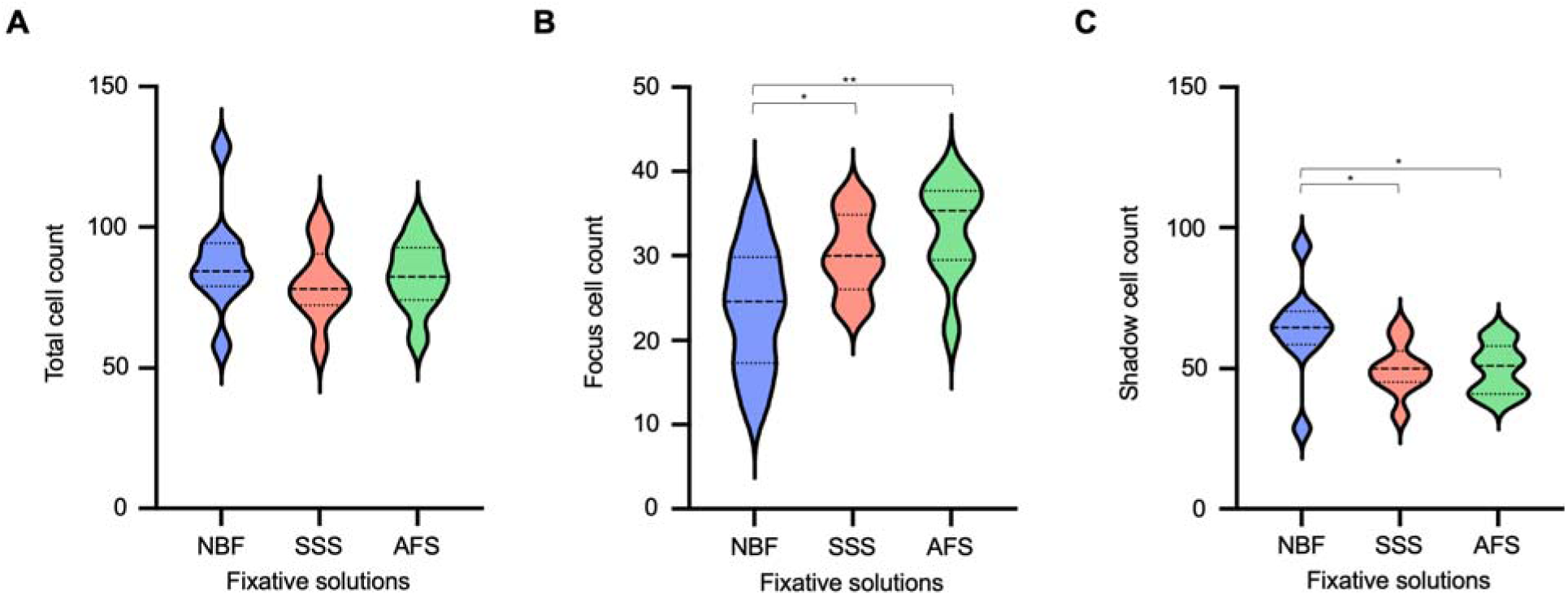
Violin plots of the cell counts in all three fixative solutions. A) Total cell count. B) In-focus cell count. C) Out-of-focus cell count. μm = micrometers; NBF = neutral-buffered formalin; SSS = saturated salt solution; AFS = alcohol-formaldehyde solution. * p < 0.05; ** p < 0.01; *** p < 0.001

#### 3.3.2 In-focus and out-of-focus cell counts

The mean number of in-focus cells per specimen was 23.81 ± 7.25 neurons in NBF-fixed specimens, 30.19 ± 4.71 neurons in SSS-fixed specimens, and 33.07 ± 5.87 neurons in AFS-fixed specimens. The number of in-focus cells significantly differed between the three fixative groups (ANOVA: F_2;24_ = 5.549; p = 0.010; η^2^_p_ = 0.316) (Figure 4B). The NBF-fixed specimens had a lower number of in-focus cells when compared with those fixed with the SSS (p = 0.035) and the AFS (p = 0.003). There was no difference between the number of in-focus cells in SSS- and AFS-fixed specimens (p = 0.320).

The mean number of out-of-focus cells per specimen was 63.74 ± 16.80 neurons in NBF-fixed specimens, 49.81 ± 9.25 neurons in SSS-fixed specimens, and 49.74 ± 8.95 neurons in AFS-fixed specimens. The out-of-focus cell numbers significantly differed between the three fixative groups (ANOVA: F_2;24_ = 3.919; p = 0.034; η^2^_p_ = 0.246) (Figure 4C). The NBF-fixed specimens had a higher number of out-of-focus cells when compared with those fixed with the SSS (p = 0.024) and the AFS (0.023). Conversely, there was no significant difference between the number of out-of-focus cells in SSS- and AFS-fixed specimens (p = 0.990).

### 3.4 Cell diameters

#### 3.4.1 Large diameters

Using the manual method, the median large diameter was 11.56 μm (range: 7.73-22.49 μm) in NBF-fixed specimens, 11.97 μm (range: 6.83-22.38 μm) in SSS-fixed specimens, and 11.96 μm (range: 5.99-24.75 μm) in AFS-fixed specimens. The manually measured large diameters were not significantly different between the cells fixed by the NBF and SSS (GLMM: *t*(804) = 0.515; p = 0.607), as well as between the cells fixed by the NBF and AFS (GLMM: *t*(804) = 0.447; p = 0.655) (Figure 5A).

**Figure 5.**
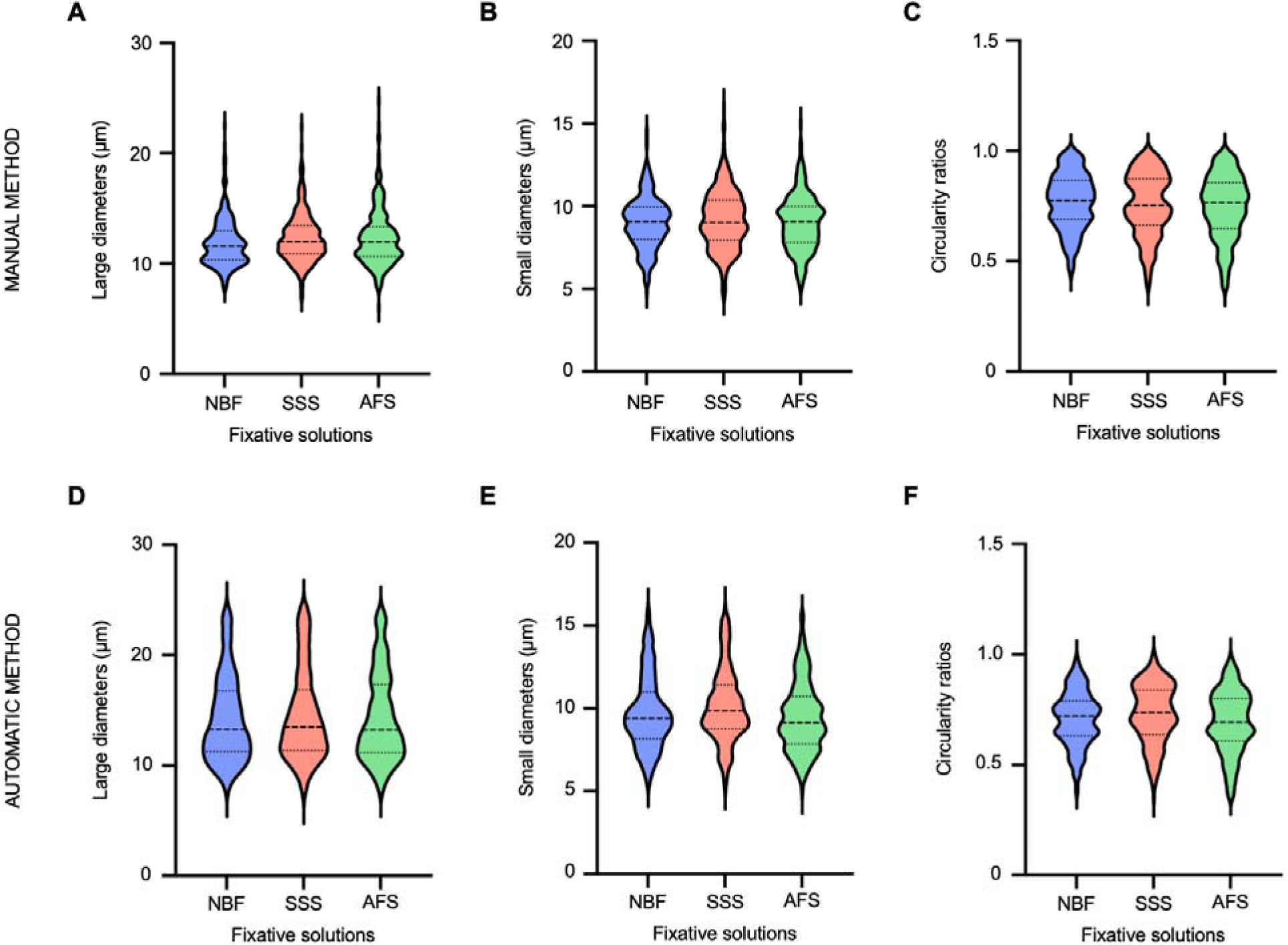
Violin plots of the cell diameters in all three fixative solutions. A) Manually measured large diameters. B) Manually measured small diameters. C) Cell circularity ratios obtained from the manual diameters. D) Automatically measured large diameters. E) Automatically measured small diameters. F) Cell circularity ratios obtained from the automatic diameters. μm = micrometers; NBF = neutral-buffered formalin; SSS = saturated salt solution; AFS = alcohol-formaldehyde solution

Using the automatic method, the median large diameter was 13.27 μm (range: 7.71-24.23 μm) in NBF-fixed specimens, 13.49 μm (range: 7.24-24.28 μm) in SSS-fixed specimens, and 13.23 μm (range: 7.81-23.75 μm) in AFS-fixed specimens (Figure 5D).

There was a significant Spearman correlation between the manually and automatically measured large diameters (r_s_ = 0.423; p < 0.001). The correlations were also significant within each fixative group (r_s_ = 0.313; p < 0.001 for NBF-fixed specimens, r_s_ = 0.453; p < 0.001 for SSS-fixed specimens, and r_s_ = 0.503; p < 0.001 for AFS-fixed specimens).

#### 3.4.2 Small diameters

Using the manual method, the median small diameters were 9.06 μm (range: 4.74-14.58 μm) in NBF-fixed specimens, 9.02 μm (range: 4.54-15.96 μm) in SSS-fixed specimens, and 9.06 μm (range: 5.05-14.95 μm) in AFS-fixed specimens. The manually measured small diameters were not significantly different between the cells fixed by the NBF and SSS (GLMM: *t*(804) = 0.701; p = 0.483) as well as between the cells fixed by the NBF and AFS (GLMM: *t*(804) = -0.590; p = 0.555) (Figure 5B).

Using the automatic method, the median small diameters were 9.41 μm (range: 5.36-15.91 μm) in NBF-fixed specimens, 9.86 μm (range: 5.14-16.10 μm) in SSS-fixed specimens, and 9.15 μm (range: 4.93-15.54 μm) in AFS-fixed specimens (Figure 5E).

There was a significant Spearman correlation between the small diameters measured manually and automatically (r_s_ = 0.469; p < 0.001). The correlations were also significant within each fixative group (r_s_ = 0.373; p < 0.001 for NBF-fixed specimens, r_s_ = 0.473; p < 0.001 for SSS-fixed specimens, and r_s_ = 0.565; p < 0.001 for AFS-fixed specimens).

#### 3.4.3 Cell circularity ratios

The manually measured cell diameters yielded a median circularity ratio of 0.77 (range: 0.44-1.00) in NBF-fixed specimens, 0.75 (range: 0.39-0.99) in SSS-fixed specimens, and 0.77 (range: 0.38-0.99) in AFS-fixed specimens. These ratios did not differ significantly between the NBF- and SSS-fixed specimens (GLMM: *t*(804) = -0.754; p = 0.451) and between the NBF- and AFS-fixed specimens (GLMM: *t*(804) = -0.301; p = 0.764) (Figure 5C).

The automatically measured cell diameters yielded a median circularity ratio of 0.72 (range: 0.38-0.99) in NBF-fixed specimens, 0.74 (range: 0.35-1.00) in SSS-fixed specimens, and 0.69 (range: 0.36-0.99) in AFS-fixed specimens (Figure 5F).

Finally, there was a significant Spearman correlation between the circularity ratios measured from the manual and automatic diameters (r_s_ = 0.444; p < 0.001). The correlations were also significant within each fixative group (r_s_ = 0.326; p < 0.001 for NBF-fixed specimens, r_s_ = 0.452; p < 0.001 for SSS-fixed specimens, and r_s_ = 0.547; p < 0.001 in AFS-fixed specimens).

## 4. Discussion

In this study, we evaluated the impact of three different fixative solutions on the quantitative analysis of neurons in the PMC of mice. Our goal was to determine whether the solutions used in human gross anatomy laboratories (i.e., the SSS and AFS) impacted the nervous tissue in a similar or idiosyncratic manner when compared to the NBF solution commonly used in brain banks. We used various quantitative variables that enabled us to assess how the solutions impacted the neuropil and neuronal size, by considering the area, number, diameters, and circularity of cells.

### 4.1 Quantitative variables

#### 4.1.1 Cell areas and number of labeled neurons

We first quantified the total cell area, by segmenting all the cells in every ROI. This showed us the overall proportion of area occupied by neurons in the same geometrical area. This variable is important to assess whether the fixation of the brain tissue and the following histology treatment had a similar or idiosyncratic impact on the tissue shrinkage or expansion in brains fixed by all three solutions. However, individually, this variable is not sufficient to conclude on the impact of the solutions on the volume of cells. The total volume could be affected by the individual volume of cells and their total number, which is why these variables were interpreted along with the total cell area.

The initial hypothesis was that NBF-fixed specimens would have a lower cell area because of the high concentration of formaldehyde in its composition, which is known to cause brain shrinkage (Mouritzen Dam, 1979). It was therefore anticipated that these specimens would contain smaller cells separated by larger areas of neuropil. Due to a previous study of our group, we also expected the AFS-fixed specimens to render larger cell areas than the other two solutions. It was observed that mounted sections of AFS-fixed mouse brains were larger than those of NBF- and SSS-fixed brains (Supplementary Figure 2) (Frigon et al., 2022a).

Our results showed that the total cell area per ROI as well as the individual cell area were not different across the three fixative solutions, invalidating our initial hypotheses. We surmise that the high concentration of alcohol in the compositions of the SSS and AFS might shrink the tissue and its cells in a similar manner as the NBF. Indeed, ethanol is also known to dehydrate tissues and diminish the volume of many organs, including the brain (Boon & Kok, 2008; Fox et al., 1985).

The results also indicated that the larger section area of AFS-fixed brains (Supplementary Figure 1) that was observed previously was not related to the size of cortical neurons, since the cell areas were not different between the three solutions. Therefore, we presumed that there might be a difference in the area of the neuropil, which would be affected independently from the cells. However, this should have impacted the number of neurons per ROI. It was not the case since the total number of labeled neurons was also not different across solutions. The observation concerning the larger area of AFS-fixed brain sections might then not be related to the impact of the fixation on the size of cortical structures, as was studied here. Instead, it might be related to the area occupied by subcortical structures and/or deep white matter, which might undergo less shrinkage after fixation with AFS and explain the larger areas of sections. This hypothesis would be consistent with the previous evidence that fixation causes a differential shrinkage and deformation in white matter as compared to gray matter (Gros et al., 2023). This, however, was not tested within the confines of this study since all ROIs are exclusively cortical. Futures studies quantifying the cells and neuropil of subcortical structures would be necessary to confirm this hypothesis.

#### 4.1.2 In-focus and out-of-focus cell counts

The purpose of distinguishing between in-focus and out-of-focus cells was to consider the three-dimensionality of the cells and the penetration of antibodies. Indeed, our hypothesis was that larger cells would generate a higher in-focus cell count, whereas smaller cells would instead generate a lower in-focus cell count. This follows the principle that larger cells have a higher probability of being cut and sampled than smaller cells (Delgado-González et al., 2020).

We initially expected AFS-fixed specimens to show more in-focus cells and less out-of-focus cells due to the observation concerning the larger area of mounted AFS-fixed sections. In other words, we expected the larger sections to hold larger cells, therefore implying a higher number of in-focus cells. Since there was no difference in cellular area or in the total neuronal count, despite the larger size of AFS-fixed brains, this hypothesis was invalidated. In fact, our findings showed that the number of in-focus cells was significantly lower and the number of out-of-focus cells was significantly higher in NBF-fixed specimens as compared to both SSS- and AFS-fixed specimens. Given that the size of the cells is similar across fixatives, we interpreted this finding as a differential penetration of antibodies in the NBF-fixed specimens, which would result in blurry labeling of cells in the middle image. In a previous study of our group (Frigon et al., 2022b), it was found that three out of nine NBF-fixed specimens showed an incomplete penetration of NeuN antibodies. We attributed this to the higher degree of protein cross-linking in these specimens, which might mask the NeuN epitopes and hinder their detection (Azumi & Battifora, 1987; Battifora & Kopinski, 1986; Boon & Kok, 2008; Boon et al., 1992; de Paula et al., 2018; Leong & Gilham, 1989; Moelans et al., 2011; Unhale et al., 2012; Wu et al., 2022). Furthermore, since the SSS and AFS contain high levels of alcohol, they might induce a more pronounced membrane permeabilization, therefore resulting in a better antibody penetration and more in-focus cells. Hence, the lack of alcohols in NBF would explain the poorer penetration of antibodies and the blurry contours of the neurons in the middle image of these specimens.

#### 4.1.3 Cell diameters and circularity ratios

Finally, we measured the cell diameters of ten randomly selected neurons per ROI, differentiating between the large and small diameters. This can be coupled with the individual cell area to assess the impact of the fixative on the cell size. Based on these diameters, we also calculated circularity ratios to characterize the neuronal morphology.

The large and small diameters were not significantly different between the three solutions, which further reinforces the notion that cell size is similarly impacted by the three fixatives. Although the difference was not significant, we observed that NBF-fixed neurons had slightly smaller large diameters than the tissue fixed by the two gross anatomy solutions. If the large diameters were indeed smaller, then the cell would have a rounder shape when compared to the more elongated form of SSS- and AFS-fixed neurons. This would point to a difference in cell morphology, which could potentially be explained by the higher level of formaldehyde in the NBF solution. However, the difference in large diameters was not statistically significant and represented less than a half-micron (11.56 vs 11.97 vs 11.96 μm), which is only 3 to 4% of the cell diameter. Consistently, the circularity ratios were also not different across solutions.

Taking all the manual results into account, nothing indicates that the quantitative variables differ across the three fixative solutions. These are promising results for the use of brains fixed by gross anatomy solutions in research.

### 4.2 Automatic software

#### 4.2.1 Total cell area

The comparison between the manual and automatic segmentation procedures showed a strong Dice coefficient (0.92) and a moderate correlation coefficient (0.532). This indicates an overall good agreement between the methods. In most cases (20 out of 27), the automatic segmentation generated higher area values than the manual segmentation. This suggests an oversegmentation by the automatic detection method. This is largely due to darker backgrounds, as shown in the upper half of Supplementary Figure 3. However, the oversegmentations in most cases are more subtle than the ones shown in the supplementary figure, usually coming from falsely segmented, darker pixels around the cells. This indicates a difference in the way to handle the inclusion or exclusion of partial volume pixels present at the borders between the cell profiles and background. Although the automatic detection software segmented slightly more pixels around the cells, the results were still similar enough to produce an overall acceptable agreement between the manual and automatic methods. Furthermore, of the seven cases in which the automatic segmentation generated lower values, three were NBF-fixed, two were SSS-fixed and two were AFS-fixed. Observations of these specimens highlighted the undersegmentation of the automatic method when the cells were very pale compared to the rest of the cells (lower half of Supplementary Figure 3). However, even with these segmentation issues in a few specimens, most cases were correctly segmented. Therefore, the automatic method is sufficiently accurate to be used on its own to segment the total cell area of fixed mouse specimens.

#### 4.2.2 Individual cell area

The automatic method for the individual cell areas was less accurate than the automatic segmentation of the total cell areas. The Dice and Spearman correlation coefficients were both lower than those observed for the total cell area (0.81 vs 0.92 and 0.50 vs 0.87, respectively). However, the Dice coefficient was still over 0.80, which is generally considered to reflect a high accuracy in pixel alignment. Moreover, the correlation coefficient was over 0.4, which reflects a strong relationship between the manual and automatic results. Although these results were generally good, a quality control (QC) of the automatic segmentations had to be performed. As shown in Supplementary Figure 1, the automatic software was unable to perform adequately in multiple cells, which were removed systematically when their area values were over twice the standard deviation. Following that procedure, 52 cells out of 810 (6.42%) were removed from the dataset. Out of these 52, there were 14 NBF-fixed cells (26.9%), 22 SSS-fixed cells (42.3%) and 16 AFS-fixed cells (30.8%). Along with a thorough visual QC of all specimens, these results indicate that SSS-fixed specimens might generate a less reliable automatic segmentation of individual cells than the other two solutions. This is probably due to their higher frequency of darker backgrounds. This qualitative observation was supported by a previous study, which reported that the background labeling with the NeuN antibody was often darker in SSS-fixed specimens (Frigon et al., 2022b).

Overall, these segmentation errors made by the automatic method did not influence the results disproportionately, apart from a minority of cells that were identified as outliers. However, we would still recommend manually correcting the segmentations to ensure the false positives and negatives are corrected before processing the data, which would be generally faster than manually segmenting every cell. Another alternative would be to apply an automatic QC step, during which the software is trained to detect segmentation failures (Sander et al., 2020).

#### 4.2.3 Cell diameters and circularity ratios

The Spearman correlation coefficient for the large and small diameters was below 0.4, which denotes a moderate agreement between the two methods. We believe the automatic measures of diameters are greatly affected by the heterogeneity of labeling intensity, as low contrast and boundary robustness are known to be problematic for accurate automatic measures (Zeng et al., 2025). Thus, the automatic method is still lacking in precision for measuring cell diameters, especially in specimens with heterogeneous labeling.

A possible explanation would be that the length of large diameters was often overestimated by the inclusion of the axon hillock and the proximal portion of the axon by the automatic software. The NeuN antibody, as was used in the current study, can stain some proximal processes (Wolf et al., 1996). However, we have noticed that this is very variable from one neuron to another, resulting in a heterogeneous population of labeled neuronal bodies with and without a partial axon. The presence of labeled axon hillocks and proximal portions of the axon would therefore falsely increase the results obtained from automatically measuring the large diameters of neurons. In contrast, a manual rater would take it into account while measuring the diameter of the neuronal body.

The overestimated large diameters using the automatic software resulted in smaller circularity ratios compared to the ones calculated from manual measures. Considering the formula of the ratios, the large diameter acts as the denominator, which means that when the denominator is higher, the resulting ratio is lower. Consequently, the automatic ratios are less reliable than the manual ratios since they are affected by the falsely higher measures of automatic large diameters.

### 4.7 Study limits

The small sample size in this study represents the main limitation to detect significant differences across fixative solutions. However, the magnitude of difference across mean values was overall small, so detecting a difference of a few microns might not be relevant for research projects looking for clear changes in cell numbers or size. A power analysis also showed that the total number of specimens needed to ensure the detection of a significant difference with a medium effect size was 269, which is simply unattainable.

Moreover, we could have assessed more quantitative variables or other antigens to compare the three fixatives. Instead, we chose to focus on the neuronal count and dimensions (area and diameters) since neuronal loss and atrophy are closely related to the pathogenesis of neurodegenerative disorders (Agnello & Ciaccio, 2022; Hammer et al., 1979; Mezzapesa et al., 2013).

Furthermore, although digital stereology could have been used to compare the quantitative variables between fixative solutions (Kristiansen & Nyengaard, 2012; Mehrabi et al., 2018), applying these methods would have required more numerous and equidistant sections of the PMC. However, it is not applicable in SSS-fixed specimens in which many sections are lost due to their difficulty of manipulation (Frigon et al., 2022b). We therefore opted for a practical option that would still allow for a comparison across groups. The ROI method allowed us to fulfill the goals of this study while still staying true to the basic sampling theory by using the different levels of thickness per slice, the multiple ROIs per specimen and the random method to select the individual cells to segment.

In addition, the ROIs were only analyzed in one region (i.e., the PMC). Consequently, the effects on each variable might differ in other regions. However, this study design was chosen explicitly to reduce the number of variables and thus ensure comparability between the three groups.

Finally, there is a concern about the relevance of these results for a human application, which is the ultimate purpose of our group’s research program. Using mouse brains to test the properties of the fixative solutions before replicating the study on human brains might seem redundant or not transferable. However, it allows us to evaluate how the brain tissue reacts to the solutions in optimal conditions without all the unavoidable confounding variables associated with human brains, such as the longer and variable postmortem delay, fixation delay, comorbidities, and biological heterogeneity of donors (Alkemade et al., 2023; Frigon et al., 2022b; Frigon et al., 2024). While the comparability of mice and humans is not the same, the study design with a single independent variable (type of fixative solution) would not be possible in humans.

## 5. Conclusion

We compared the effects of fixative solutions from gross anatomy laboratories on neurons in delimited regions of interest in the primary motor cortex. The cell area, total number of neurons and cell diameters were not different across the three fixatives tested, reflecting their comparability in histology measures. Since this work focused on mouse tissue, future studies applying a similar methodology on human tissue will determine its suitability for quantitative analyses. Overall, these results are promising for neuroscientists interested in using brains from anatomy laboratories for quantitative research on cortical neurons.

## Acknowledgements

The authors would like to thank the funding resources and the staff at the Animal Care facilities of the University of Québec in Trois-Rivières.

