## Supplementary material for "Neuronal quantification in the primary motor cortex of mouse brains fixed with solutions from human gross anatomy laboratories"

Corresponding author:

Amy Gérin-Lajoie

Université du Québec à Trois-Rivières

Acknowledgements: The authors would like to thank the funding resources and the staff at the Animal Care facilities of the University of Québec in Trois-Rivières.

Funding: This work was funded by the NSERC grant to JM. AGL was supported by the NSERC and FRQNT.

Conflict of interest: The authors declare that the research was conducted in the absence of any commercial or financial relationships that could be construed as a potential conflict of interest.

Word count: 6 787

Number of supplementary figures and tables: respectively 3 and 1

### **Supplementary figures**

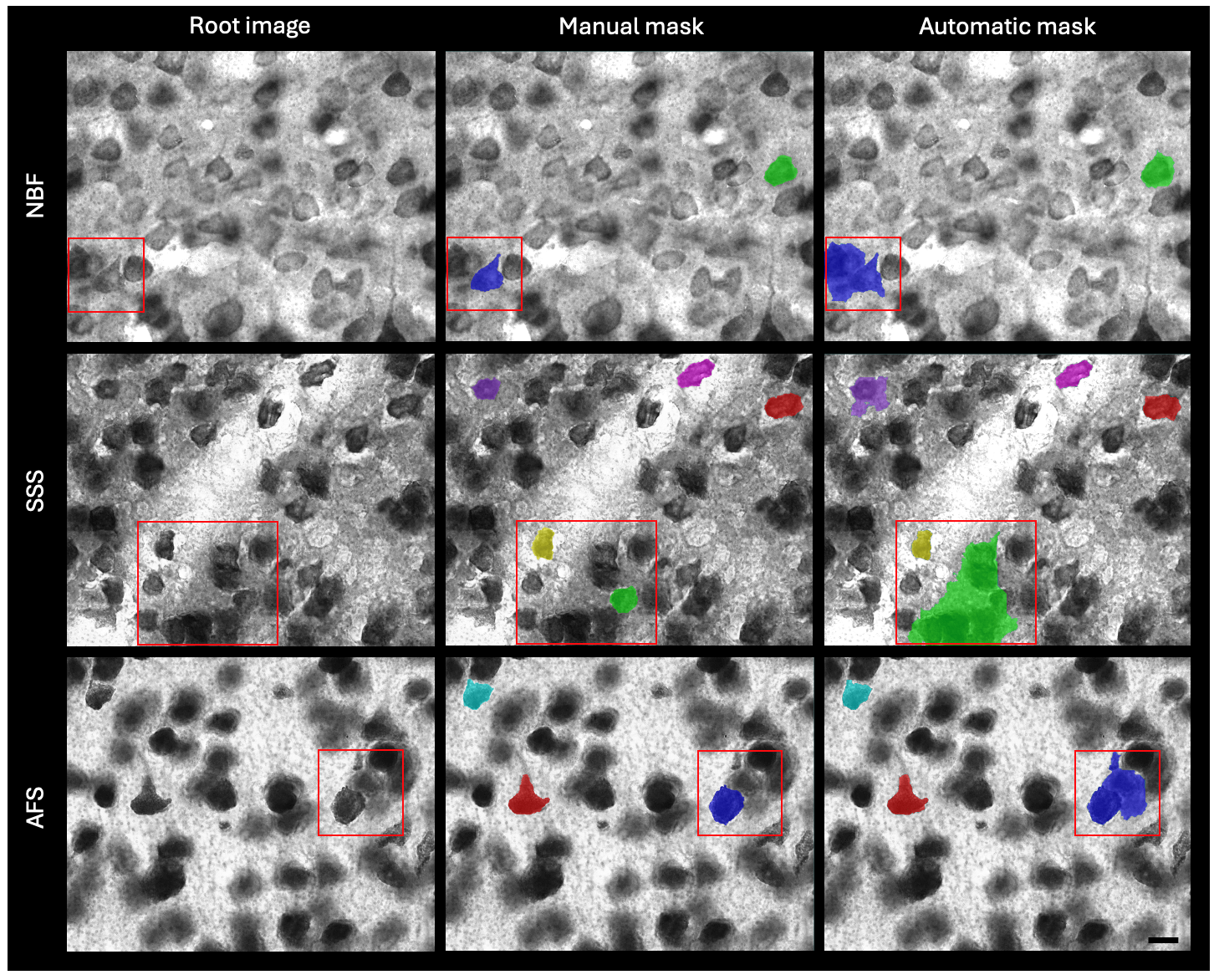

Supplementary Figure 1. - **Outliers in the automatic segmentation of individual cells**. Red squares identify the specific cells that were removed from further statistical analyses, before segmentation (left column), after manual segmentation (middle column) and after automatic segmentation (right column). Each row shows one specimen fixed by each of the solution. NBF = neutral-buffered formalin; SSS = saturated salt solution; AFS = alcohol-formaldehyde solution. Scalebar = 10 μm.

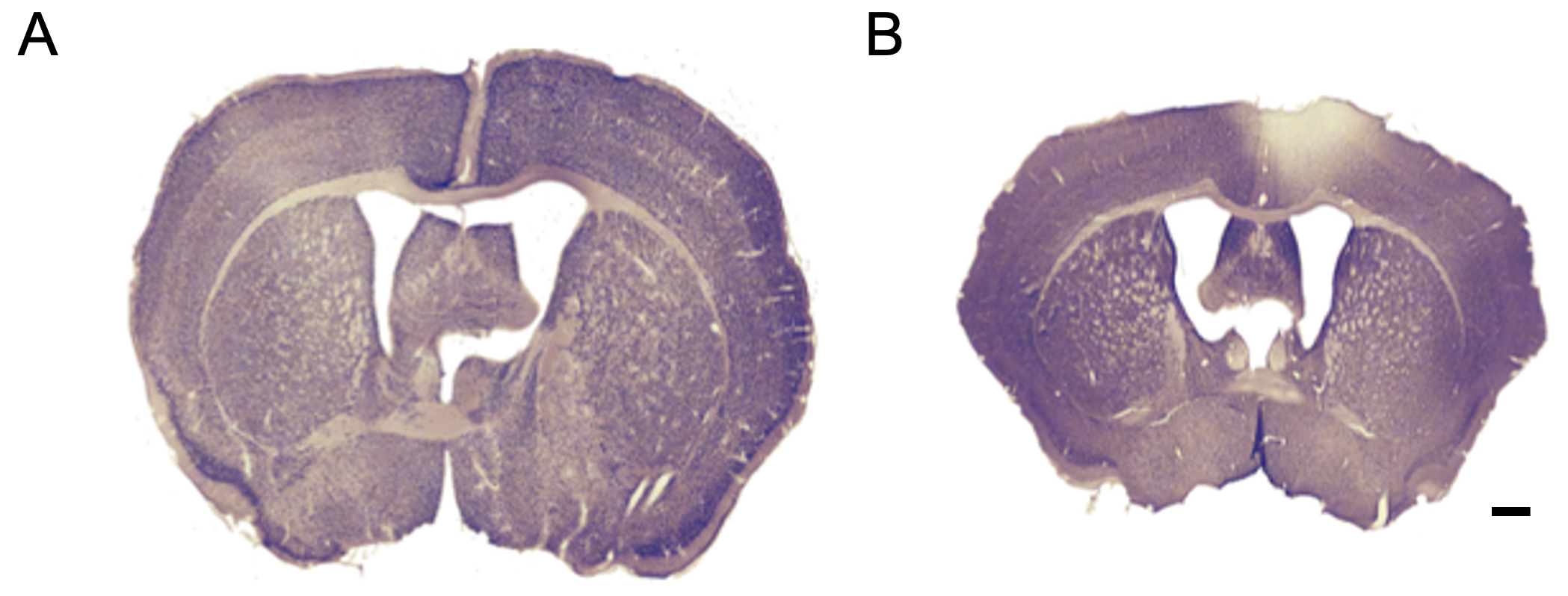

Supplementary Figure 2. - **Mounted sections of fixed mouse brains**. A) AFS-fixed section. B) NBF-fixed section. AFS = alcohol-formaldehyde solution; NBF = neutral-buffered formalin. Scalebar = 500 μm.

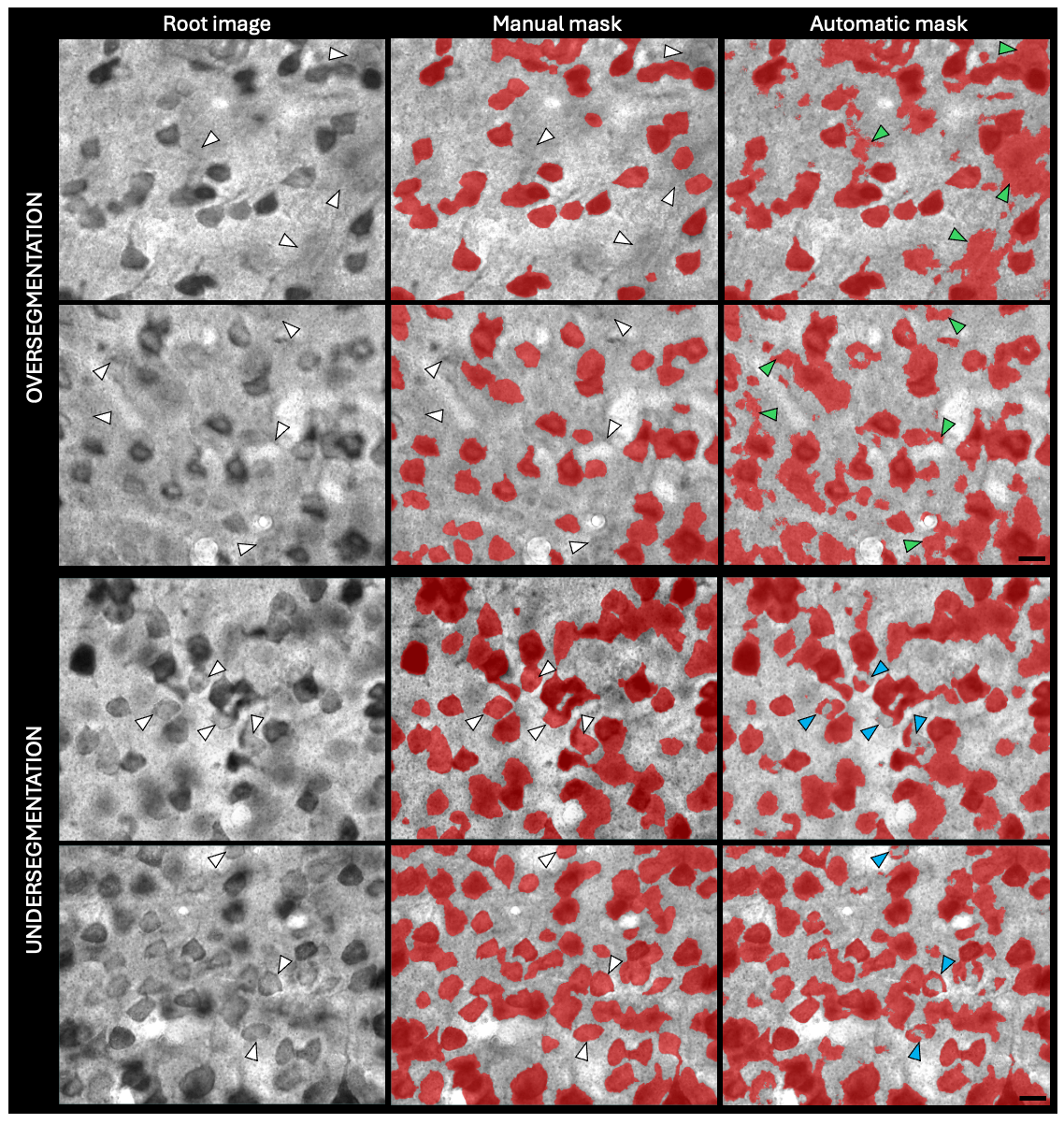

Supplementary Figure 3. - **Segmentation errors of the automatic method for the total cell area**. White arrowheads point to the regions affected by the segmentation errors in the root images and manual masks. Green and blue arrowheads respectively point to the oversegmentation and undersegmentation errors on the automatic masks. All specimens shown here are NBF-fixed. NBF = neutral-buffered formalin. Scalebar = 10 μm.

### **Supplementary Tables**

|  |  | **With outliers** | **Without outliers**  **(3x SD)** | **Without outliers**  **(2x SD)** |
| --- | --- | --- | --- | --- |
| **Individual cell area** (μm^2^) | Mean | 114.58 | 108.76 | 103.55 |
|  | SD | 72.52 | 56.39 | 47.78 |
|  | Median | 93.05 | 91.71 | 89.94 |
|  | Range | 32.92 - 756.87 | 32.92 - 323.48 | 32.92 - 259.47 |
| **Large diameters** (μm) | Mean | 14.86 | 14.67 | 14.36 |
|  | SD | 4.74 | 4.37 | 3.97 |
|  | Median | 13.52 | 13.47 | 13.32 |
|  | Range | 7.24 - 41.65 | 7.24 - 28.93 | 7.24 - 24.28 |
| **Small diameters** (μm) | Mean | 10.26 | 10.09 | 9.79 |
|  | SD | 2.96 | 2.60 | 2.16 |
|  | Median | 9.63 | 9.58 | 9.50 |
|  | Range | 4.93 - 28.84 | 4.93 - 18.99 | 4.93 - 16.10 |
| **Circularity ratios** | Mean | 0.71 | 0.71 | 0.71 |
|  | SD | 0.13 | 0.13 | 0.13 |
|  | Median | 0.72 | 0.72 | 0.72 |
|  | Range | 0.35 - 1.00 | 0.35 - 1.00 | 0.35 - 1.00 |

Supplementary Table 1. - **Descriptive statistics of the automatic measures for individual cells before and after outliers**. All solutions were combined. 3x SD = three times the standard deviation; 2x SD = two times the standard deviation; SD = standard deviation; μm = microns
